# Functional primary human 3D skeletal muscle models enable exercise and metabolic research

**DOI:** 10.64898/2026.06.29.735246

**Authors:** Simon I. Dreher, Robin Schöler, Katharina S. Zorn, Jens Martin, Jana Kühnle, Kevin Elsner, Imme Behle, Thomas Goj, Lara Ruoff, Kolja Leffek, Alessia Moruzzi, Peter Loskill, Andre Tomalka, Tobias Siebert, Andreas L. Birkenfeld, Andreas Peter, Cora Weigert

**Affiliations:** Institute for Clinical Chemistry and Pathobiochemistry, Department for Diagnostic Laboratory Medicine, University Hospital Tübingen, 72076 Tübingen, Germany; Department for Microphysiological Systems, Institute of Biomedical Engineering, Faculty of Medicine, Eberhard Karls University Tübingen, 72074 Tübingen, Germany; NMI Natural and Medical Sciences Institute at the University of Tübingen, 72770 Reutlingen, Germany; Motion and Exercise Science, University of Stuttgart, 70569 Stuttgart, Germany; Internal Medicine IV, Department of Diabetology, Endocrinology and Nephrology, University Hospital Tübingen, Otfried-Müller-Str. 10, 72076 Tübingen, Germany; Institute for Diabetes Research and Metabolic Diseases of Helmholtz Munich at the University of Tübingen, 72076 Tübingen, Germany; German Center for Diabetes Research (DZD), 72076 Tübingen, Germany

**Keywords:** Diabetes, Exercise, Human skeletal muscle organoids, Insulin, Metformin, Testosterone, TGFbeta

## Abstract

Human skeletal muscle is the principal site of insulin-stimulated glucose disposal and a major mediator of exercise-induced metabolic benefits, yet human models that preserve metabolic and exercise responsiveness remain limited. We generated primary human skeletal muscle organoids from donor-derived CD56+ myoblasts using a collagen-based extracellular matrix and serum-free IGF1-guided differentiation. The organoids formed aligned contractile tissues containing oxidative and glycolytic fiber type-like myotubes, displayed enhanced mitochondrial respiration, insulin-stimulated glucose uptake, and reproducible force generation. Electrical pulse stimulation induced AMPK activation, increased glucose utilization and lactate production, and upregulated canonical exercise-responsive genes including *NR4A3* and *PPARGC1A*. Notably, transcriptional responses to in vitro exercise overlapped with acute exercise responses observed in skeletal muscle biopsies from the same donors. The organoids further detected functional impairments of skeletal muscle performance induced by TGF-β1 and metformin and increased speed generation by testosterone treatment. These findings establish a donor-specific human skeletal muscle platform that recapitulates key features of insulin action and exercise adaptation and may enable mechanistic studies of skeletal muscle metabolism, exercise responsiveness, and therapeutic interventions relevant to diabetes.

**Graphical Abstract:** 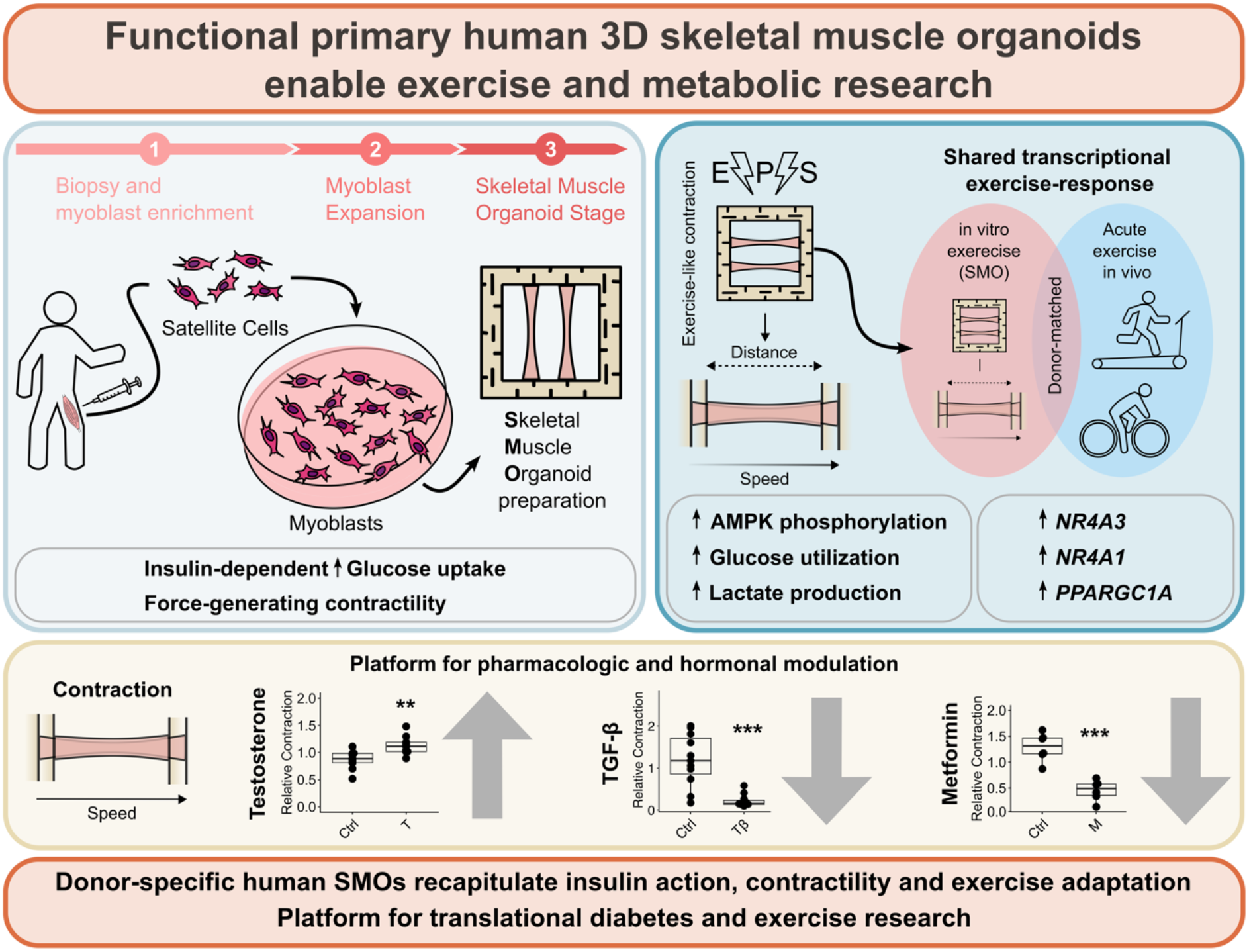

**Article highlights:** We generated primary human skeletal muscle organoids under serum-free IGF1-guided conditions to reproduce key metabolic and exercise-responsive features of skeletal muscle.

The organoids were insulin-responsive, displayed enhanced mitochondrial function and force-generating contractility, reproduced hallmark molecular and metabolic responses to exercise, overlapping with exercise responses observed in the same donors in vivo.

The organoids were suitable to detect functional alterations after treatment with endogenous hormones and cytokines and diabetes medication

This platform provides a human donor-specific system for studying skeletal muscle mechanisms underlying insulin sensitivity, exercise benefits, and therapeutic responses relevant to diabetes and metabolic disease.

## 1 Introduction

Enhanced physical activity is one of the most effective lifestyle strategies to preserve insulin sensitivity in healthy individuals and improve glycemic control in people with prediabetes or type 2 diabetes [1; 2]. Skeletal muscle is central to these benefits. It accounts for the majority of insulin-stimulated glucose disposal in humans and adapts to exercise through increased oxidative capacity, capillarization, mitochondrial content, contractile remodeling, and enhanced insulin-dependent glucose uptake [3–7]. Beyond its role as a metabolic sink, skeletal muscle also acts as an endocrine and paracrine organ, releasing myokines and myometabokines that coordinate local and systemic adaptations to exercise [8–10]. However, the molecular mechanisms linking contracting human skeletal muscle to whole-body metabolic health remain incompletely understood.

Progress in this area requires experimental models that capture physiologically relevant human muscle function while allowing controlled mechanistic interrogation. Human intervention studies provide the highest translational relevance but are limited by restricted access to tissue biopsies, limited suitability for systematic compound or metabolite screening, and interindividual variability. Mouse models enable integrated organ-level studies, but species-specific differences in anatomy, metabolism, molecular regulation, and housing physiology can limit direct translation to humans [11; 12]. Human in vitro systems therefore provide an important complementary strategy, particularly when designed to reduce animal use in accordance with the 3Rs principle and to support organ-on-chip or multicellular co-culture approaches [13; 14].

Primary human myoblast-derived myotubes are widely used to study skeletal muscle biology. Established protocols can generate long, multinucleated myotubes that reproduce selected features of mature myofibers [15]. However, conventional two-dimensional differentiation systems often remain limited by incomplete structural, contractile, and metabolic maturation [16]. We recently developed a serum-free, IGF1-guided differentiation protocol for primary human myoblasts that promotes a more mature metabolic and contractile phenotype in two-dimensional culture [17]. IGF1-supported myotubes showed increased diameter, multinucleation, physiological contraction upon electrical pulse stimulation, a proteomic shift toward sarcomeric and oxidative muscle proteins, elevated mitochondrial respiration, increased GLUT4 abundance, and enhanced insulin-dependent glucose uptake [17]. These findings established a metabolically competent human muscle model, but its two-dimensional architecture remains intrinsically non-physiological.

Three-dimensional human skeletal muscle models offer an opportunity to better reproduce the anisotropic architecture, cell–matrix interactions, and contractile mechanics of native muscle. Conventional spherical organoids are poorly suited for modeling skeletal muscle exercise physiology because they do not recapitulate the aligned, longitudinal organization required for force generation and contraction-based adaptation [18]. Accordingly, engineered skeletal muscle organoids and tissues are commonly generated as elongated constructs, often using fibrin/thrombin-based matrices, and have been successfully applied to study contraction, loading, hypertrophy, quiescent satellite-like states, glucose uptake, and force production [19–22]. Nevertheless, many current 3D skeletal muscle systems primarily emphasize structural or contractile maturation, whereas robust metabolic maturation, insulin responsiveness, and human exercise-like adaptation remain less well established.

The extracellular matrix is another key determinant of physiological relevance. Native skeletal muscle is embedded in a collagen-and laminin-rich matrix that provides mechanical support, regulates fiber stability, is essential for force transmission, shapes mechanotransduction, and contributes to the satellite cell niche [23–25]. Collagen-based 3D culture systems may therefore provide a more tissue-relevant environment than fibrin-dominated constructs, although previous collagen-based approaches have often relied on cell lines or culture conditions that are not optimized for metabolic disease research [26]. In addition, many 3D differentiation strategies still use insulin-containing media [27–29], limiting their suitability for investigating insulin sensitivity, insulin resistance, and diabetes-relevant skeletal muscle physiology.

Thus, a central unmet need remains. Current human skeletal muscle models reproduce selected structural or contractile features of muscle tissue, but inadequately combine primary human donor origin, physiological 3D architecture, metabolic maturation, insulin responsiveness, and acute exercise-like adaptability.

Here, we hypothesized that combining primary human myoblasts, collagen-based 3D tissue engineering, and serum-free IGF1-guided differentiation would generate metabolically mature, contractile, and exercise-responsive human skeletal muscle organoids. We therefore aimed to develop a primary human collagen-based skeletal muscle organoid model, adapt IGF1-guided differentiation to this 3D system, and characterize its structural, metabolic, contractile, and acute exercise-responsive phenotype. To strengthen physiological relevance, we compared in vitro acute exercise responses with acute exercise performed by the same human donors. Finally, we evaluated whether this novel human skeletal muscle organoid (SMO) captures functional responses to testosterone, TGF-β, and metformin, thereby testing its utility for studying anabolic, catabolic, and metabolic modulation in a human-relevant setting.

## 2 Methods

### 2.1 Donors

Primary human myoblasts were obtained from muscle biopsies as described previously [17]. In brief, muscle biopsies were taken from the lateral portion of the vastus lateralis of the quadriceps femoris after local anesthesia (2% Scandicaine; AstraZeneca, Germany) under sterile conditions using a fine-needle punch biopsy technique (Peter Pflugbeil GmbH, Germany). Biopsy donors were participants of previous intervention studies [30; 31] and only baseline biopsies were used for in vitro experiments. Biopsies from a total of 36 donors were used in this study, 17 females, 19 males, age 30±8 years, BMI 29±5. Most donors (BMI 31±4) suffered from overweight with a high risk to develop diabetes and 7 donors with a BMI below 25 (BMI 22±1) were included. All participants gave written informed consent, and the study protocols were approved by the ethics committee of the University of Tübingen and in accordance with the Declaration of Helsinki. For experiments comparing in vitro and in vivo exercise responses, skeletal muscle organoids were generated from the same donors who underwent acute exercise testing.

### 2.2 Exercise intervention

The in vivo 8-weeks-exercise intervention was performed as described previously [32]. Participants performed one hour of supervised endurance training three times per week, consisting of 30min of cycling on an ergometer and 30min of walking on a treadmill. Peak VO2 was defined as the mean VO2 over the last 20s before the cessation of exercise and was assessed by metabolic gas analysis. The training intensity was individually set at 80% of the VO2peak determined before the intervention. Training intensity was controlled by heart rate based on predetermined 80% of the VO2peak and individually set. Biopsies used for in vivo transcriptome data were taken during a baseline visit 8 days before and 60min after the first ergometer exercise bout.

### 2.3 Myoblast isolation

Primary human satellite cell-derived myoblasts were isolated from vastus lateralis muscle biopsies as described previously [17]. CD56-positive myoblasts were enriched by magnetic cell sorting and expanded before use in two-dimensional or three-dimensional culture as described in [33]. In brief, human satellite cells were released by collagenase digestion and seeded on 15cm dishes coated with GelTrex™ (Thermo Fisher Scientific, Cat#A1413302, USA). After two rounds of proliferation in cloning medium (α-MEM:Ham’s F-12 (1:1), 20% (v/v) FBS, 1% (v/v) chicken extract, 2mM L-glutamine, 100units/ml penicillin, 100µg/ml streptomycin, 0.5µg/ml amphotericin B), CD56-positive myoblasts were enriched (>90%, 96±3% over all donors) using MACS microbeads and LS columns (Miltenyi Biotec, Germany), according to the manufacturer’s protocol, with a 30min incubation. They were then stored in the gaseous phase of liquid nitrogen. Cell culture surfaces were prepared with a non-gelling thin-layer GelTrex™ coating. Myoblasts (passage 3 after isolation, passage 1 after enrichment) were proliferated in cloning media until 90% confluency.

### 2.4 2D culture and treatment

Primary human myoblasts were differentiated in serum-free differentiation medium supplemented with IGF1 as described previously [17]. In brief, myotube differentiation was induced on day 0 and maintained for 7-8 days in fusion media (α-MEM containing 5.5mM glucose, 2mM L-glutamine, 50µM palmitate, 50µM oleate (complexed to BSA with a final BSA concentration of 1.6mg/ml in medium), 100µM carnitine) with or without supplementation of IGF1 (human recombinant IGF1, Sigma-Aldrich, Cat#I3769, Germany). Medium was changed three times per week and 48h before harvest. Mycoplasma-free culture conditions and cells are regularly controlled for using the MycoAlert Mycoplasma Detection Kit (Lonza, Switzerland). For IGF1 withdrawal experiments, myotubes were differentiated in the presence of IGF1 and IGF1 was removed at defined time points before harvest. For IGF1 dose-response experiments, myoblasts were differentiated for 8 days in serum-free differentiation medium containing 100, 50, 25, or 10ng/ml IGF1. For analysis of insulin-stimulated AKT phosphorylation, IGF1 was removed from the medium for 48h before stimulation with 10nM insulin.

### 2.5 3D culture and treatment

Longitudinal skeletal muscle organoids were generated from expanded primary human CD56-positive myoblasts. Myoblasts (500,000 cells per SMO) were embedded in an extracellular matrix consisting of 10% TeloCol-6 (Advanced BioMatrix Inc., Cat#5225, USA), 20% Geltrex (Thermo Fisher Scientific, Cat#A1413302, USA). Two organoids were cast in one PDMS mold with two channels leading to an inserted nylon frame (Cerex, PBN-II 30, 4.0osy, 136gsm, USA). After 24h, SMOs anchored to the nylon frame to maintain longitudinal tension and tissue alignment, were removed from the PDMS mold and transferred into 6-well plates with differentiation medium. Organoids were differentiated for 7-8 days in serum-free differentiation medium supplemented with 25ng/ml IGF1 before further analysis. For analysis of insulin-stimulated glucose uptake, SMOs were fasted for 48h before stimulation with 100 or 1000nM insulin. For the testosterone response testing, in accordance with previous primary human myotube cell-culture experiments, cells were treated throughout the differentiation with 100nM testosterone (Sigma-Aldrich, Cat#86500, Germany) or the solvent control (0.006% ethanol) [32]. For the TGF-β response testing, cells were treated for 48h before harvest with 1ng/ml TGF-β1 (RCD Systems, Cat# 7754-BH005, USA), 10µM TGF-β signaling inhibitor SB431542 (Sigma-Aldrich, Cat#S4317, Germany), 1ng/ml TGF-β1 and 10µM SB431542 or the solvent control (DMSO 1:26 and 4mM HCl/0.1% BSA 1:50). For the metformin response testing, cells were treated for 48h before harvest with 78µM or 776µM metformin hydrochloride (Cayman Chemical, Cat#13118, USA). Further controls were left untreated. To induce contraction, the “Move” protocol was applied in all cases.

### 2.6 Contraction and in vitro exercise

Skeletal muscle cells in 2D and SMOs were exercised using electrical-pulse-stimulation (EPS) to induce controlled contraction. Controls contained electrodes for the same time period. Cells and organoids were cultured in compatible 6-well plates (Falcon, Corning, USA). After complete differentiation at day 7-8, EPS was applied using C-Pace EP (IonOptix, USA). Three stimulation protocols were used: “Move,” consisting of one controlled contraction every 2s for 20h (0.5Hz, 4ms, 12V), “Intense,” consisting of tetanic contraction for 2s followed by 2s relaxation for 2h (10Hz/0Hz, 4ms, 12V), and “Fast,” consisting of two contractions per second for 2h (2Hz, 4ms, 12V). To assess general contractility, EPS was applied to cells and SMOs for 10min using “Move”. Videos of 15s at 60fps (frames per second) per group and donor were taken in random spots of each well for 2D culture or at the center of the SMOs using an Axiovert 40C (Carl Zeiss Microscopy, Germany) with Flexacam C3 Camera (Leica, Germany). Videos were analyzed using a motion-tracking algorithm [34]. For the analysis, a block width of 16 pixels, as well as a delay of two frames and a maximum shift of 20 pixels was used. For each SMO, a region of interest (750px x 120px or 1616µm x 256µm, at the center of SMOs) was selected. The algorithm captures velocity of movement [µm/s] over time, allowing the measurement of contraction speed by averaging the peaks measured over 10s. For quantification of the contraction distance, the area under the curve (AUC) was calculated over 10s of measurement.

### 2.7 Force measurement

SMOs were activated by calcium diffusion (pCa 4.5) in the presence of ATP, following protocols described in detail in Tomalka et al. (2017) [35]. Each SMO was mounted between a high-speed length controller (322C-I, Aurora Scientific, Canada) and a force transducer (403A, Aurora Scientific, Canada) using T-shaped aluminum clips attached to both ends. Before each experiment, the SMO was stretched until passive force became detectable and all slack was removed. This length was defined as the resting length (*Lrest*), corresponding to a passive baseline force of 5–10 µN. To determine active force-generating capacity, SMOs were activated isometrically at *Lrest*, and maximum active force was quantified as the peak force attained during contraction. Subsequently, passive mechanical properties were assessed by stretching SMOs from *Lrest* by 10%, 20%, and 40% of their resting length over a constant duration of 2 s. Passive force was quantified as the peak force reached at the end of each stretch, allowing assessment of length-dependent passive force development.

### 2.8 RNA analysis

Transcriptomic analysis of skeletal muscle biopsies [32] and qPCR analysis [17] was performed as described previously. For qPCR, myotubes in 2D and SMOs were homogenized in RLT lysis buffer (Qiagen, Cat#79216, Germany). Proteinase K digestion (Qiagen, Cat#19134, Germany) and DNAse I solution (Qiagen, Cat#79254, Germany) were used. For generating cDNA, First Strand cDNA Synthesis Kit (Roche, Cat#04897030001, Switzerland) was performed. RNA targets were quantified using SYBR Green Universal Master Mix (Thermo Scientific, Cat#4309155, USA), primers from Qiagen (Germany) of TBP (QT00000721), RPS28 (QT02310203), MYH1 (QT01671005), MYH2 (QT00082495), MYH7 (QT00000602), PPARGC1A (QT00095578), GLUT4 (QT00097902), NR4A3 (QT01669941) and NR4A1 (QT00095515) with a LightCycler480 (Roche, Switzerland).

### 2.9 Western blot

Protein abundance and phosphorylation were analyzed by Western blot as described before [17; 36]. Summarized, 2D cultures and SMOs were lysed using RIPA buffer (25mM Tris pH 7.6, 150mM NaCl, 3.5mM SDS, 12.1mM sodiumdeoxycholate, 1% (v/v) Triton X100) containing cOmplete EDTA free protease inhibitor (Roche Diagnostics, Cat#11697498001, Switzerland) and phosphatase inhibitor (1mM NaF, 0.5mM sodium pyrophosphate, 1mM β glycerophosphate, 1mM sodium orthovanadate). Protein content was quantified utilizing a BCA assay kit (Thermo Scientific, Cat#23225, USA). 10-15µg protein, denaturated in Laemmli buffer were loaded onto a gradient polyacrylamide gel (5-15%). Semidry transfer was performed using a Nitrocellulose membrane. For blocking, NET-G buffer (NaCl (0.15M), Tris-EDTA (5mM), Tris-HCl (50mM), gelatine 0.25% (w/v), pH∼7.4) was utilized. Primary antibody incubation was overnight at 4°C, while secondary antibodies were incubated for 2h at RT. Immunosignals were analyzed with an Odyssey detection machine (LI-COR Biosciences, USA). The analyzed targets included MYH1/2 (Sigma-Aldrich, Cat#M4276, Germany; Clone MY-32), MYH7 (Sigma-Aldrich, Cat#M8421, Germany; Clone NOQ7.5.4D), GLUT4 (Invitrogen, Cat#PA5-23052, USA), phosphorylated AKT (Cell Signaling Technology, Cat#9271L, USA), total AKT (BD Biosciences, Cat#610860, USA), total AMPK (Merck Millipore/Upstate, Cat#07-363, Germany), phosphorylated AMPK (Cell Signaling Technology, Cat#2535, USA) and GAPDH (Abcam, Cat#ab8245, United Kingdom).

### 2.10 Histology

SMOs were analyzed by histology and immunostaining to assess myotube formation, longitudinal alignment, and fiber-type marker expression. SMOs were fixed in 4% PFA and permeabilized overnight at RT (Permeabilization Solution containing stabilizer and 0.09% azide (Miltenyi Biotec, Cat#130-126-719, Germany). Incubation with primary and secondary antibodies was performed for 24-48h each at 37°C in Antibody Staining Solution (Miltenyi Biotec, Cat#130-126-719, Germany). After dehydration in increasing EtOH/2% Tween20 solution (50%-70%-100% EtOH), stained SMOs were cleared using Clearing Solution (Miltenyi Biotec, Cat#130-126-719, Germany). Proteins were stained with monoclonal anti-Myosin, MHC-fast, Clone MY-32 (1:1000, Sigma-Aldrich, Cat.#M4276, Germany) and monoclonal anti-Myosin, MHC-slow, Clone NOQ7.5.4D (1:1000, Sigma-Aldrich, Cat.#M8421, Germany), Goat anti-Rabbit IgG (H+L) Highly Cross-Adsorbed Secondary Antibody, Alexa Fluor™ Plus 488 (Invitrogen, Cat.#A32731, USA) and Donkey anti-Mouse IgG (H+L) Highly Cross-Adsorbed Secondary Antibody, Alexa Fluor™ 568 (Invitrogen, Cat.#A10037, USA) secondary antibody as well as DAPI (1:500, Invitrogen, USA) and imaged using the ApoTome System (Carl Zeiss Microscopy, Germany).

### 2.11 High-resolution respirometry

Mitochondrial respiration was assessed using Seahorse extracellular flux analysis. SMOs were differentiated in 6-well plates with or without IGF1 for 7 days. For the Seahorse experiment, an XFe24 sensor cartridge and XFe24 Seahorse Islet Capture microplate were utilized (Agilent, Cat#103518-100, USA). The sensor cartridge was calibrated according to the manufacturer. The XFe24 Seahorse Islet Capture microplate was coated with a cell/tissue adhesive (Corning® Cell-Tak™, Cat#354240, USA). A volume of 0.5µL (c=1.4µg/µL, pH=4.0) was added to the wells. Then, 10µL Seahorse assay medium (pH=7.4) was placed on top of the Cell-Tak solution. Seahorse assay medium consisted of 97% (v/v) Seahorse medium (Agilent, Cat#103575-100, USA), 1% (v/v) 1M glucose solution (Agilent, Cat#103577-100, USA), 1% (v/v) 100mM pyruvate (Agilent, Cat#103578-100, USA), and 1% (v/v) 200mM glutamine (Agilent, Cat#103579-100, USA). Further incubation at RT for 20min led to an optimal adherence effect. The Seahorse sensor cartridge was loaded in Port1 with oligomycin (10µM), Port2 with FCCP (20µM), and finally Port3 with a mixture of rotenone (5µM) and antimycin A (5µM). SMOs on day 7 of differentiation were washed twice with Seahorse assay medium. Then, individual organoids were detached from the frame with forceps and placed in the Cell-Tak prepared XFe24 Seahorse Islet Capture microplate after aspirating the remaining Seahorse assay medium. Carefully, 100µL Seahorse assay medium was added and the plate was centrifuged at 200xg at RT for 1min with speeding and breaking at level 2 (5810R Eppendorf centrifuge). The volume of all wells was filled to 500µL in total with Seahorse assay medium. After calibration, the XFe24 Seahorse Islet Capture microplate with the attached SMOs in Seahorse assay medium was placed in the instrument. Settings were set to 3min mixing, 2min incubation and 3min measurement per condition. After the Seahorse experiment, protein determination was performed with a BCA assay kit (Thermo Scientific, Cat#23225, USA). OCR values of the Seahorse experiment were normalized to total protein content of the particular SMO.

### 2.12 Glucose and Lactate measurement

SMOs were differentiated according to the standard protocol for 7 days in fusion medium. The SMOs were then either exercised with the “Move” protocol as described above or left unexercised as a control for the same period of time. Immediately afterwards, the supernatants were harvested and centrifuged at 500g for 5min at RT. Due to evaporation during overnight electrical pulse stimulation, supernatant volumes were measured to allow volume correction of the data. Lactate and glucose concentrations were measured using the Atellica Solution CH Analyzer (Siemens Healthineers AG, Germany).

### 2.13 Glucose uptake

Glucose uptake was assessed as described previously [17] and after in vitro exercise stimulation using the “Intense” protocol. Glucose uptake was determined using the Glucose Uptake-Glo™ Assay (Promega, Cat#J1343, USA) following the protocol provided by the manufacturer. SMOs were cultured for 7 days before being exercised for 2h in 6-well plates as described above, followed by 24h of fasting. Incubation with 100nM or 1000nM insulin (Roche Diagnostics, Switzerland) was performed for 1h at 37°C, followed by 1h-incubation at room temperature after adding 10mM 2-deoxyglucose. Luminescence was measured using the GloMax® Multi Detection System (Promega, USA) after 1h-incubation at room temperature with the provided detection reagent.

### 2.14 Statistics

Differences between individual groups were assessed using one-way ANOVA with Fisheŕs LSD or Welch post hoc test depending on normal distribution and Bonferroni correction for multiple comparisons when appropriate. Graphs were made using the R packages ‘ggplot2’ (v3.3.2) and figures assembled using InkScape (v1.0). Data points represent individual donors, box plots show the median, interquartile range, and minimum–maximum values. Where appropriate, data were normalized per donor. Statistical analysis was performed in R4.5.1 [37].

## 3 Results

### 3.1 Functional myotube maturation at 25ng/ml IGF1

Exogenous IGF1 (100ng/ml) in combination with a serum-free medium was key for achieving a mature and contractile phenotype of myotubes. Before transferring the system to a more advanced 3D organoid culture, we validated optimal timing and dose of IGF1 administration. The presence of IGF1 in the medium for the complete differentiation period of 8 days (D8) or removal 2 days before harvest on day 6 (D6) had no measurable impact on myotube contractility or myosin heavy chains as differentiation markers on both transcript and protein level (Fig.S 1 A-F). Only RNA expression of *PGC1a*, *GLUT4* and GLUT4 protein was reduced after 2 days without IGF1 (Fig.S 1 G-I). Removing IGF1 already on day 4 or 2 (D4, D2) strongly decreased almost all analyzed parameters (Fig.S 1). The comparison of IGF1 concentrations of 100, 50, 25, and 10ng/ml during 8 days of differentiation revealed no significant differences in the same set of markers, however, inter-donor variability increased at 10ng/ml (Fig.S 2). To achieve a pronounced effect of insulin (10nM) on AKT phosphorylation, removal of IGF1 for 48h was required independent of the added concentration of IGF1 (Fig.S 3). Removing IGF1 for 24h already achieved a clearly visible trend of insulin-induced AKT phosphorylation.

### 3.2 Generation of 3D primary human skeletal muscle organoids

Based on the knowledge gained in 2D culture, we subsequently used 25ng/ml IGF1 in the differentiation medium and removed IGF1 for 24 or 48h if needed for assessment of insulin effects. We generated longitudinally shaped 3D skeletal muscle organoids (SMOs) also starting from human CD56+ myoblasts enriched by magnetic cell sorting of human vastus lateralis-derived myoblasts (Fig. 1). Expanded myoblasts were cast into a physiology-inspired ECM containing type 1 collagen, laminin, heparan sulfate and proteoglycans. Two SMOs were cast inside a PDMS mold anchoring the SMOs between a nylon frame supporting tissue tension maintaining the longitudinal shape similarly to fascia connecting the skeletal muscle to bone. Myoblasts inside the SMOs were subsequently cultured for 7 days in serum-free differentiation medium to allow complete functional maturation before further analysis (Fig. 1).

**Fig. 1.**
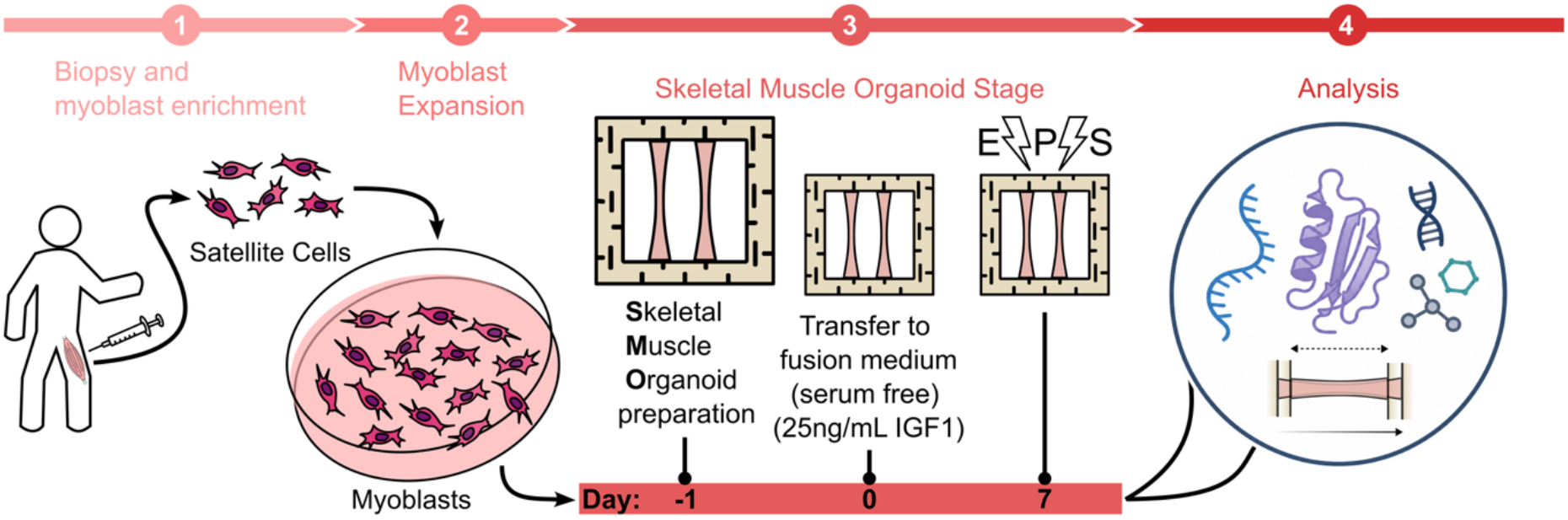
Generation of primary human skeletal muscle organoids. Schematic representation of the workflow starting from primary human satellite cell isolation, myoblast enrichment, to SMO generation, culture and analysis.

### 3.3 Baseline characterization of skeletal muscle organoids

Similarly to what we previously demonstrated for 2D myotube culture, SMOs differentiated in the presence of IGF1 contain both MYH1/2 positive fast type 2 glycolytic as well as MYH7 positive slow type 1 oxidative myotubes (Fig. 2 A). Fully differentiated SMOs on day 7 maintained their shape throughout culture (10mm longitudinally and 0.51±0.13mm diameter). In contrast to 2D culture, differentiating myotubes linearly aligned along the ECM and longitudinal axis of the SMO. In line with previous data in 2D culture, the presence of IGF1 during differentiation led to elevated mitochondrial respiration compared to SMOs differentiated without IGF1 (Fig. 2 B). Insulin-dependent glucose uptake can be observed in SMOs as well (Fig. 2 C). When applying electrical pulse stimulation (EPS) to SMOs in culture, pronounced contractility was observed when cultured with IGF1 in the media, and this contractility was induced repeatedly and reproducibly from day 5 of culture over several days (Fig. 2 D, E). While contractions were more pronounced when SMOs were fully differentiated on day 7 compared to day 5 (0h vs 0h, 2h vs 2h, 7h vs 7h), an initial adaptation to EPS within the first 2h after the first application of electrical pulses (0h vs 7h respectively) enabled full contractility further on, regardless of the differentiation status (Fig. 2 F). This adaptation of the contractile apparatus aligned with force measurements performed on our SMOs, where both passive and active force were elevated after 2h of EPS at 0.5Hz by around 20-80µN (Fig. 2 G-L). Active forces reached values of up to 600µN and 470µN for the EPS and control conditions, respectively (Fig. 2 J,L). Taken together, we successfully implemented a physiological, insulin responsive and contractile 3D skeletal muscle organoid based on primary CD56+ myoblasts from human donors.

**Fig. 2.**
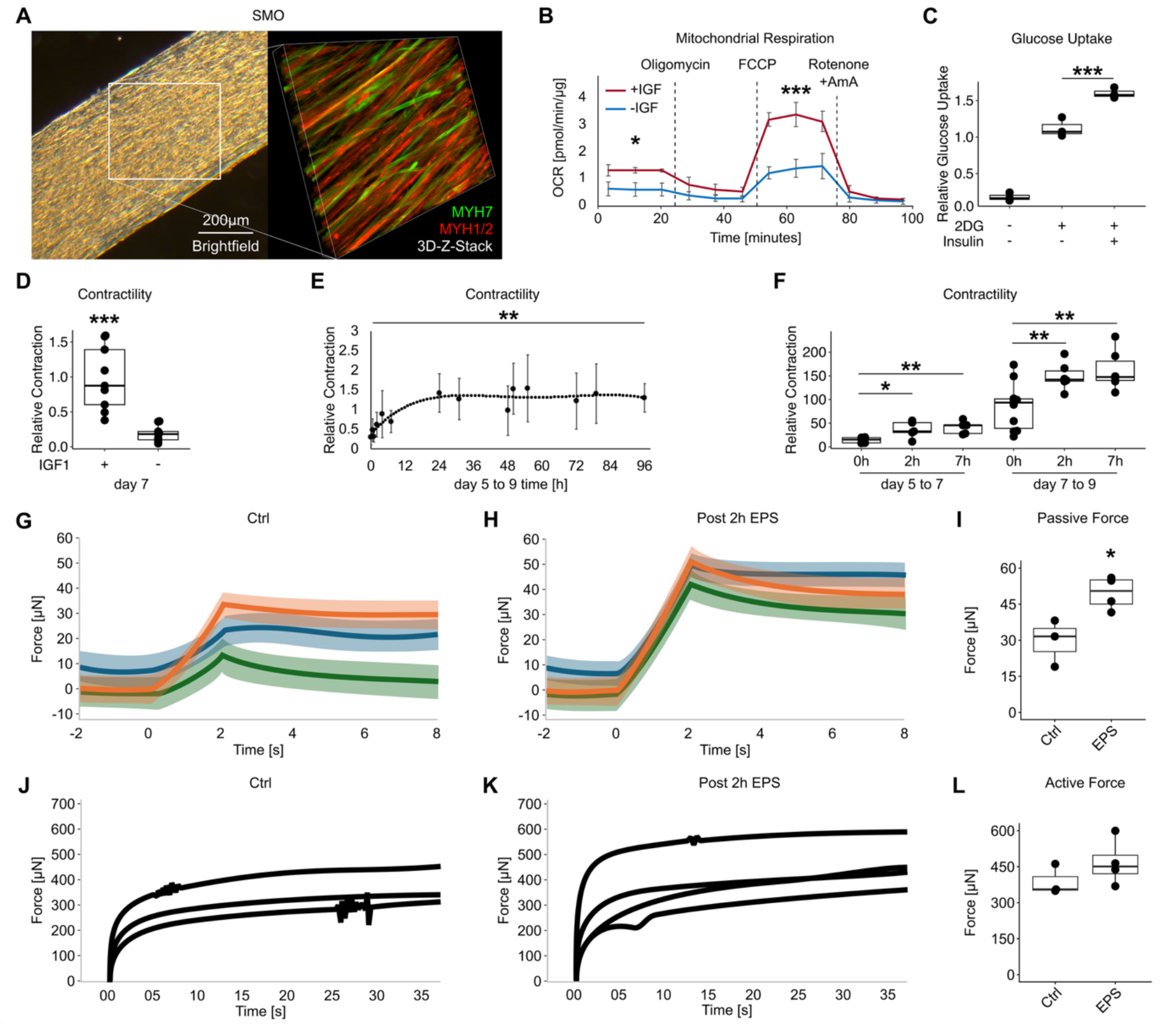
Baseline characterization of skeletal muscle organoids. SMOs were generated from primary human myoblasts and differentiated for 7 days before analysis. (A) Brightfield overview and immunostaining of MYH1 positive fast type glycolytic and MYH7 positive slow type oxidative myotubes. (B) High resolution respirometry of SMOs differentiated in the presence or absence of IGF1. (C) Glucose uptake in SMOs stimulated by 100nM Insulin after 48h without IGF1. (D) Contractility in SMOs differentiated in the presence or absence of IGF1. (E) Repeated measurement of contractility between day 5 and 9 of differentiation. (F) Contractility measured in the first 7h starting on day 5 or 7 of differentiation. (G-I) SMOs were subjected to passive stretches of increasing amplitudes (10 [green], 20 [blue], 40% [orange]) over a constant duration of 2 s in the control condition (Ctrl; G) and 2 h after EPS (H) at 0.5 Hz. Colored traces represent force responses at different stretch amplitudes; shaded areas indicate variability across samples. Passive force increased with stretch amplitude and was higher following EPS (H). Peak passive force values are summarized in panel I. (J-L) Representative black traces show active isometric force recordings in control (J) and post-EPS (K) SMOs following activation by calcium diffusion (pCa 4.5) in the presence of ATP. Maximum force-generating capacity was determined from peak isometric force. Summary data are presented in panel L. Contractility is represented by contraction distance. Statistical significance was determined by one-way ANOVA with Fisheŕs LSD or Welch post hoc test depending on normal distribution and Bonferroni correction for multiple comparisons when appropriate, n=3-10 individual donors, *p<0.05, **p<0.01, ***p<0.001.

### 3.4 Skeletal muscle organoid acute in vitro exercise response

Next, we characterized acute exercise responses of SMOs applying different EPS modalities (Fig. 3). We defined 3 exercise modalities, with the moderate “Move”, SMOs performed a controlled contraction every 2s for 20h, “Intense” induced tetanic contraction for 2s followed by a 2s-relaxation phase repeatedly for 2h and in the “Fast” protocol SMOs contracted twice per second for 2h (Fig. 3 A). Increased phosphorylation of AMPK (pAMPK/AMPK) was observed directly after “Intense”, “Fast” and 1h after “Intense” exercise, as well as after “Move” by trend despite also observing slightly elevated total AMPK levels in response to EPS (Fig. 3 B). We then analyzed the transcriptional response in SMOs and compared it to the transcriptional response in skeletal muscle biopsies of the same human donors after an acute bout of exercise [32]. Typical acute exercise response genes *NR4A3* and *PPARGC1A* (PGC1a) were strongly upregulated after exercise in vivo as well as in SMOs after “Intense” EPS, while *NR4A3* was also elevated in response to the “Fast” protocol as well as 1h after “Intense and “Fast” (Fig. 3 C-H). Interestingly, unlike in skeletal muscle biopsies, both “Move” and “Intense” slightly elevated expression of differentiation markers *MYH1/2* in SMOs while *MYH7* remained unchanged by exercise in vitro and in vivo (Fig.S 4). Increased glucose utilization and lactate production were observed in the supernatant of SMOs following 20h of “Move” (Fig. 3 I). Application of 2h “Intense” exercise increased glucose uptake to a similar extent as insulin-induced glucose uptake in control conditions (Fig. 3 N). Altogether, response of SMOs to in vitro exercise in different modalities reflects key aspects of the physiological response to exercise in vivo.

**Fig. 3.**
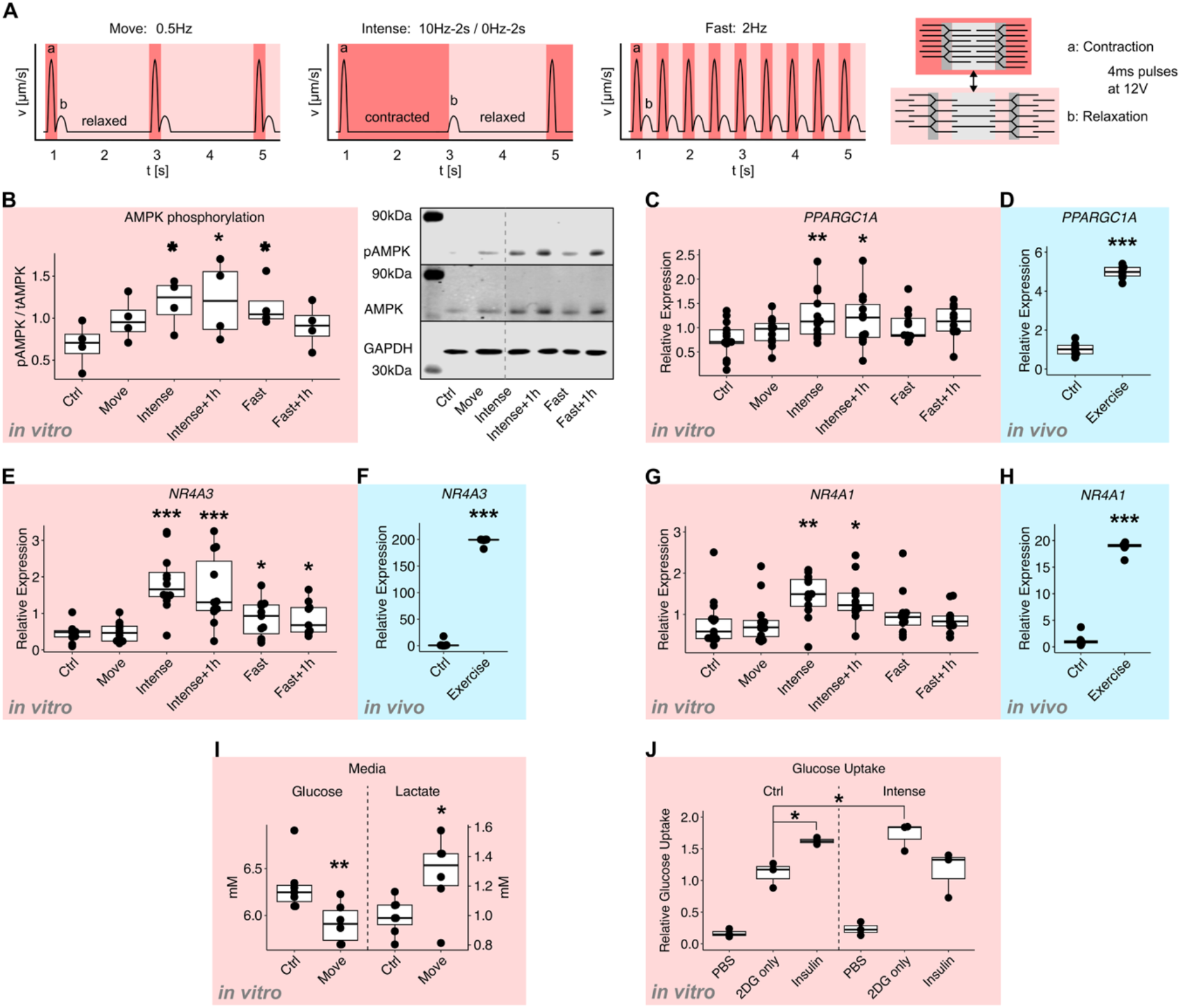
Skeletal muscle organoid in vitro exercise. SMOs were generated from primary human myoblasts and differentiated for 7 days before analysis. (A) Schematic representation of in vitro exercise protocols “Move” 0.5Hz for 20h, “Intense” 10Hz-2s followed by 0Hz-2s for 2h intermittent and “Fast” 2Hz for 2h with 4ms pulses at 12V. (B) Thr-172 phosphorylation of AMPK, total AMPK and GAPDH as loading control were measured by Western blot. Quantification of n=4 individual donors and a representative blot are shown. RNA expression of *PPARGC1A* (*PGC1a*) (C, D), *NR4A3* (E, F) and *NR4A1* (G, H) were measured in SMOs directly or 1h (+1h) after in vitro exercise at indicated protocols compared to unexercised controls (Ctrl) (C, E, G) and in skeletal muscle biopsies of the same human donors SMOs were generated from before (Ctrl) and 1h after exercise (Exercise) (D, F, H). (I) Glucose and Lactate measured in SMO culture medium after exercise with the “Move” protocol or without (Ctrl). (J) Glucose uptake in SMOs without (Ctrl) or after 2h of exercise using the “Intense” protocol with or without 1000nM Insulin after 48h without IGF1. Statistical significance was determined by one-way ANOVA with Fisheŕs LSD or Welch post hoc test depending on normal distribution and Bonferroni correction for multiple comparisons when appropriate, n=3-12 individual donors, *p<0.05, **p<0.01, ***p<0.001.

### 3.5 Modulators of human skeletal muscle contractility

One major functional response now enabled by these novel SMOs is contractility. As a proof of concept, we analyzed this functional outcome when treating SMOs with substances expected to affect the performance of skeletal muscles (Fig. 4 A). Testosterone is a potent substance to enhance muscle mass and strength, leading to improved contraction kinetics and power generation in human muscle fibers [38]. When treating our SMOs generated from both female and male donors with testosterone during differentiation, specifically contraction speed but not distance was increased independent of donor’s sex (Fig. 4 B-C, Fig.S 5 A-B). Local TGF-β signaling in skeletal muscle was identified to be elevated in individuals showing an impaired response to exercise and was shown to reduce myotube differentiation and insulin signaling in traditional 2D cultures [36]. When treating SMOs for 48h with TGF-β1, both contraction distance and speed were drastically diminished, which was rescued by co-treatment with a TGF-β signaling inhibitor (Fig. 4 D-E). Finally, potential adverse effects of metformin on the improvement in cardiorespiratory fitness, exercise performance and metabolic control after exercise interventions have been discussed for years [39]. When treating SMOs with metformin, both contraction distance and speed were impaired in a dose-dependent manner, with strongly diminished contractile function at the high dose (Fig. 4 F-G). To conclude, the insulin-responsieve and contractile SMOs open a new way to study diabetes and exercise-related questions in a skeletal muscle model with potentially donor-specific properties. They can also be used to study the functional impact of substances on human skeletal muscle functionality that cannot be studied in humans directly.

**Fig. 4.**
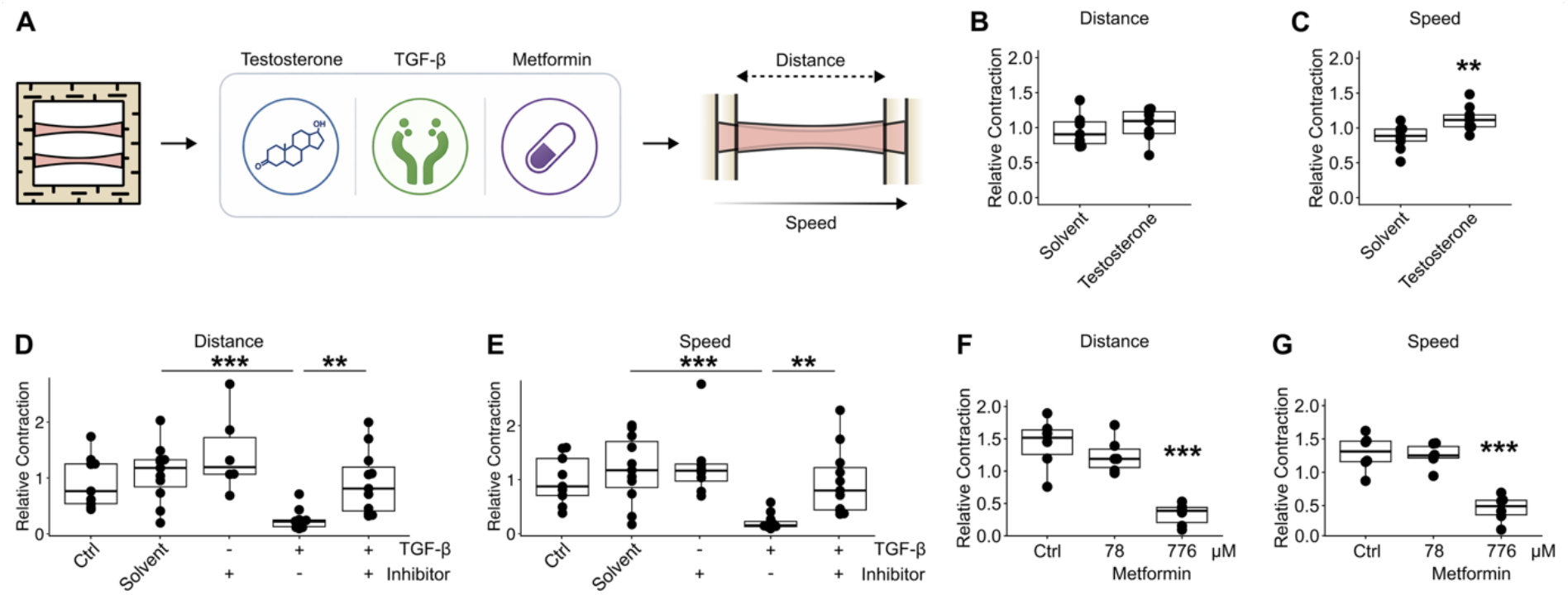
Modulators of human skeletal muscle contractility. SMOs were generated from primary human myoblasts and differentiated for 7 days before analysis. (A) Schematic representation of treatment and analysis of SMO contraction speed and distance. Contraction distance and speed of SMOs stimulated with the “Move” protocol were measured (B-C) after treating SMOs during 7 days of differentiation with or without 100nM testosterone, (C-E) for 48h before harvest with 1ng/ml TGF-β1, 10µM TGF-β signaling inhibitor SB431542 or both or (F-G) for 48h before harvest with 78µM or 776µM metformin. Statistical significance was determined by one-way ANOVA with Fisheŕs LSD or Welch post hoc test depending on normal distribution and Bonferroni correction for multiple comparisons when appropriate, n=6-11 individual donors, *p<0.05, **p<0.01, ***p<0.001.

## 4 Discussion

In this study, we established a primary human 3D skeletal muscle organoid (SMO) model that combines aligned tissue architecture, serum-free metabolic maturation, insulin responsiveness, reproducible contractility, and acute exercise responsiveness. Primary CD56+ myoblasts differentiated within a collagen-and laminin-containing extracellular matrix (ECM) into longitudinally aligned SMOs containing both MYH7+ oxidative/type 1-like and MYH1/2+ glycolytic/type 2-like myotubes. Due to serum-free IGF1 guided differentiation, the tissues showed enhanced mitochondrial respiration, insulin-stimulated glucose uptake, and reproducible electrically induced contractions with quantifiable force and contraction kinetics. Together, these features position the model as a donor-specific and mechanically active human skeletal muscle platform for studying exercise biology, metabolism, and pharmacological modulation.

The current SMO system is conceptually related to previously described engineered human skeletal muscle platforms but occupies a distinct niche. Primary human “myobundles” established that engineered human muscle can form aligned, contractile tissues responsive to electrical and pharmacological stimulation [27], while subsequent iPSC-derived systems demonstrated progressive maturation and force generation in 3D culture [40]. However, many engineered muscle systems prioritize structural maturation or neuromuscular modeling rather than metabolic functionality. In contrast, the present model was specifically designed to preserve insulin responsiveness and enable controlled investigation of exercise and metabolic adaptation.

A central distinction is the use of a collagen-, laminin-, and proteoglycan-rich ECM rather than fibrin-dominant hydrogels commonly used in engineered muscle systems. Skeletal muscle ECM regulates force transmission, tissue organization, and muscle stem cell behavior [23; 24], while matrix elasticity strongly influences myogenic differentiation and self-renewal [41]. The robust alignment, sustained contractility, and repeated EPS responsiveness observed in the SMOs likely emerge in part from recreating a more physiological biomechanical environment. This may also explain the improved structural stability during prolonged contraction compared with conventional 2D myotube systems, which frequently delaminate during extensive stimulation.

An equally important feature is the serum-free IGF1-guided differentiation strategy. Chronic insulin exposure is known to impair insulin signaling and induce insulin resistance-like phenotypes in cultured murine skeletal muscle cells [42]. Skeletal muscle insulin resistance is characterized by impaired insulin-stimulated glucose uptake, altered intracellular signaling, and dysregulated GLUT4 trafficking [43]. By separating anabolic support for differentiation from continuous supraphysiological insulin exposure, the current protocol preserves insulin-dependent glucose uptake and insulin-responsive AKT signaling similarly to what we demonstrated in 2D culture [17]. This provides an important advantage for diabetes-related research questions and drug-exercise interaction studies. In addition, the absence of serum reduces interference from undefined cytokines, hormones, and growth factors, allowing controlled addition of single substances and more targeted mechanistic investigation. Because the SMOs are generated from primary human donor-derived cells, the platform in future studies enables investigation of donor-specific variables including sex, age, obesity, training status, and diabetes, essential for precision-medicine approaches. The present platform may therefore occupy a unique niche among engineered skeletal muscle systems by specifically prioritizing metabolic functionality and insulin responsiveness rather than solely structural maturation or contractility.

The SMOs also provide a controlled platform for studying exercise-like skeletal muscle function. EPS has long been used as an in vitro surrogate for exercise, and cultured human myotubes exposed to EPS reproduce some contraction-associated metabolic and secretory responses [44; 45]. In the present model, distinct EPS protocols induced exercise-associated signaling and metabolic responses, including AMPK phosphorylation, elevated glucose utilization, lactate release, and increased *NR4A3* and *PPARGC1A* expression. These findings are consistent with transcriptomic studies identifying conserved exercise-responsive gene programs in human skeletal muscle [46]. Previous observations suggest that primary human myotubes preserve some aspects of donor metabolic phenotype after isolation and culture [47]. The model therefore provides an opportunity to investigate inter-individual variability in exercise adaptation under highly controlled conditions.

One notable finding was the rapid enhancement of contractile performance after the initial EPS bout, despite prior morphological maturation of the SMOs. Acute contraction induces intracellular Ca²⁺ transients that activate Ca²⁺/calmodulin-dependent signaling pathways involved in skeletal muscle plasticity [48]. Prior activation can also induce post-activation potentiation through increased Ca²⁺ sensitivity of the contractile apparatus and phosphorylation of myosin regulatory light chains [49; 50]. The observed increase in active and passive force may therefore reflect rapid functional priming of the contractile machinery. These findings suggest that engineered human skeletal muscle retains dynamic acute plasticity resembling early exercise adaptation in vivo.

The pharmacological perturbation experiments further demonstrate the translational potential of the system. Testosterone is well established to increase muscle size and strength in humans [38; 51]. In the SMOs, testosterone increased contraction speed without increasing contraction distance, suggesting effects on excitation-contraction coupling, calcium handling, or contractile kinetics rather than force amplitude alone. The significant decrease in contraction distance and speed following TGF-β1 treatment highlights the harmful effects of chronic TGF-β1 activation on muscle health and performance, and its part in impaired exercise adaptation [30; 36; 52]. Metformin also dose-dependently impaired contractile function. Given reports that metformin can blunt selected adaptations to aerobic exercise training [39], these findings suggest that the SMOs may provide a human muscle-specific platform to dissect drug-exercise interactions mechanistically. Importantly, the model links molecular perturbations directly to measurable functional outcomes.

The ability to quantify force generation and contraction kinetics gives the platform substantial translational potential. Generally, skeletal muscle toxicity is a clinically important adverse effect of several therapeutic classes, including statins, glucocorticoids, and chemotherapeutics [53–55]. Donor-specific SMOs could therefore enable individualized testing of anabolic therapies, anti-fibrotic compounds, cachexia interventions, metabolic drugs, and exercise-drug interactions. The structural stability of the tissues during repeated EPS also makes the system attractive for studying contraction-induced secretory signaling. Longitudinal sampling of conditioned media after defined EPS protocols may therefore allow donor-specific analysis of exercise-induced secretomes.

An additional conceptual strength of the current SMOs is the ability to study muscle-intrinsic adaptation independently of systemic influences. In vivo exercise responses emerge from complex interactions between skeletal muscle, circulation, endocrine signaling, immune cells, and the nervous system. By isolating the muscle compartment, the present model enables controlled investigation of contraction-dependent adaptation before introducing additional biological complexity. Future incorporation of motor neurons, endothelial cells, immune cells, fibro-adipogenic progenitors, or adipocytes may further extend physiological relevance while preserving the current strengths of the model.

Some limitations should be acknowledged. The SMOs are not innervated and therefore cannot fully recapitulate neuromuscular junction maturation, physiological motor unit recruitment, or neuron-dependent fiber-type specification. In addition, the model lacks vascular perfusion, endocrine dynamics, and immune interactions. The EPS protocols should therefore be interpreted as operational in vitro exercise paradigms rather than direct equivalents of endurance or resistance exercise. Potentially due to these factors and despite displaying insulin stimulated and in vitro exercise elevated glucose uptake, in SMOs, no additive effect of in vitro exercise and insulin stimulation were observed at the measured timepoints. Exercise responses in vivo appear more pronounced and are likely more complex because they integrate neural activation, blood flow, hormonal signaling, substrate delivery, and systemic inflammation. Nevertheless, the reductionist design is also a major advantage for mechanistic studies focused specifically on muscle-intrinsic adaptation.

### 4.1 Conclusion

In conclusion, this study establishes a primary human 3D skeletal muscle organoid model that combines serum-free metabolic maturation, insulin responsiveness, aligned contractility, force measurement, and exercise-like adaptation. Relative to existing engineered muscle systems, the model fills an important niche as a donor-specific and metabolically responsive model for human exercise and diabetes research. The ability to directly quantify functional adaptation after pharmacological or exercise-like stimulation provides a translational framework for studying personalized exercise biology, drug-exercise interactions, metabolic disease, and skeletal muscle-targeted therapeutics.

## 5 Acknowledgments

The authors thank all biopsy donors.

## 8 Author Contributions

S.I.D, P.L, T.S and C.W conceptualized the study. S.I.D, R.S, K.S.Z, J.M, J.K, K.E, I.B, L.R, K.L, A.M, A.T and C.W devised the methodology. S.I.D, T.G, T.S conducted the formal analysis and supervised the project. S.I.D curated the data. S.I.D wrote the original draft of the manuscript. S.I.D, R.S, K.S.Z, J.M, J.K., K.E, I.B, T.G, L.R, K.L, A.M, P.L, A.T, T.S, A.L.B, A.P and C.W reviewed and edited the manuscript. S.I.D and C.W supervised the study. S.I.D, T.S, A.L.B, A.P and C.W acquired funding. S.I.D and C.W are the guarantors of this work and, as such, had full access to all the data in the study and takes responsibility for the integrity of the data and the accuracy of the data analysis.

---

**Fig. S1.**
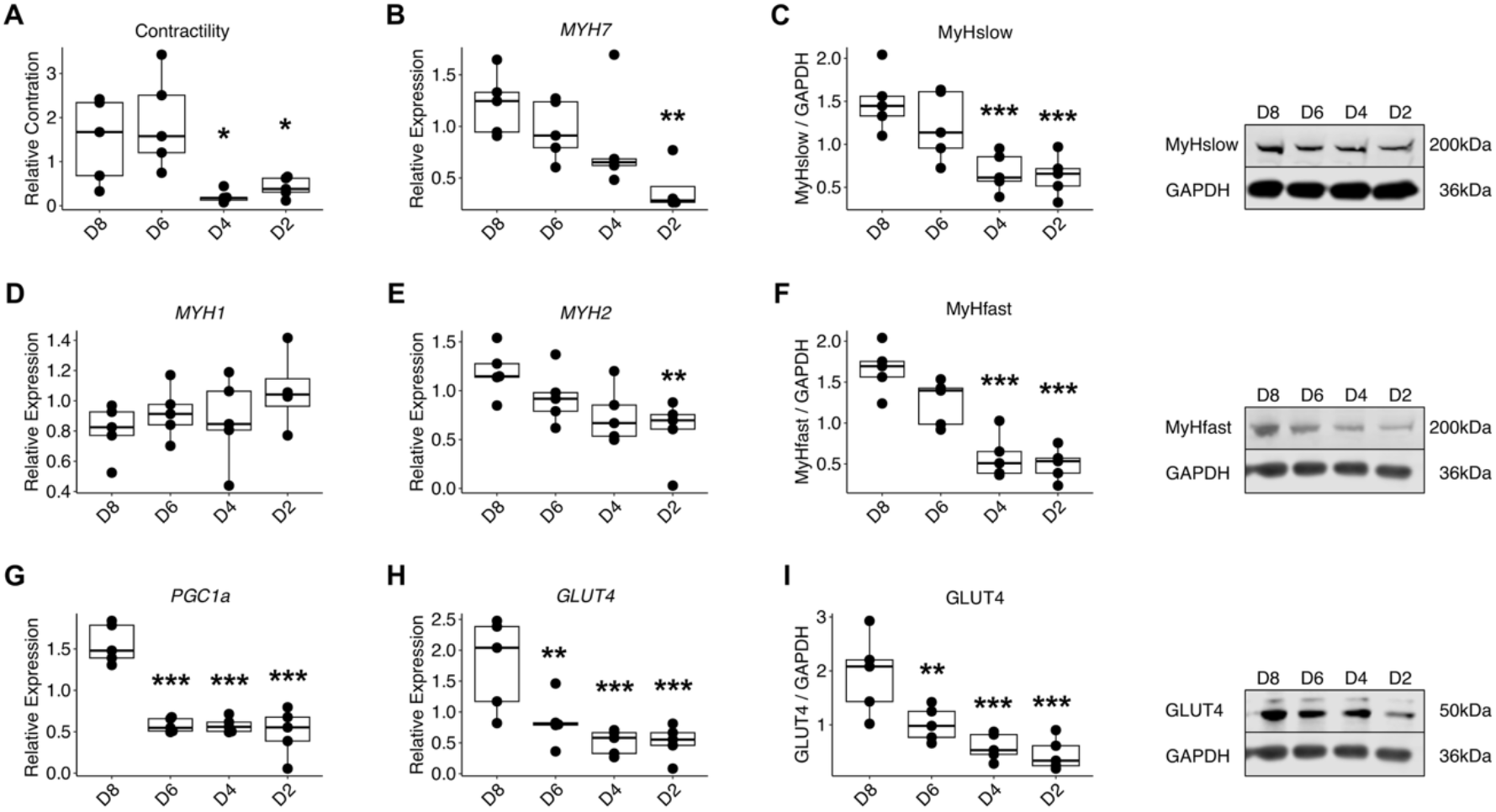
Time-resolved necessity for IGF1 during myotube differentiation. Primary human myotubes were differentiated for 8 days in the presence of insulin-like growth factor1 (IGF1) to analyze contraction and differentiation. IGF1 (100ng/ml) was removed on day 2 (D2), 4 (D4), 6 (D6) of differentiation or kept until harvest on day 8 (D8). (A) Contractility represented by contraction distance. RNA expression of differentiation markers *MYH7* (B), *MYH1* (D), *MYH2* (E), *PGC1a* (*PPARGC1A*) (G) and *GLUT4* (H) were measured on day 8 via qPCR. Protein levels of MyHslow (MYH7) (C), MyHfast (MYH1/2) (F) and GLUT4 (I) were measured by Western blot. Representative blots are shown. Statistical significance was determined by one-way ANOVA with Bonferroni correction for multiple comparisons, n=4-5 individual donors, *p<0.05, **p<0.01, ***p<0.001 vs D8.

**Fig. S2.**
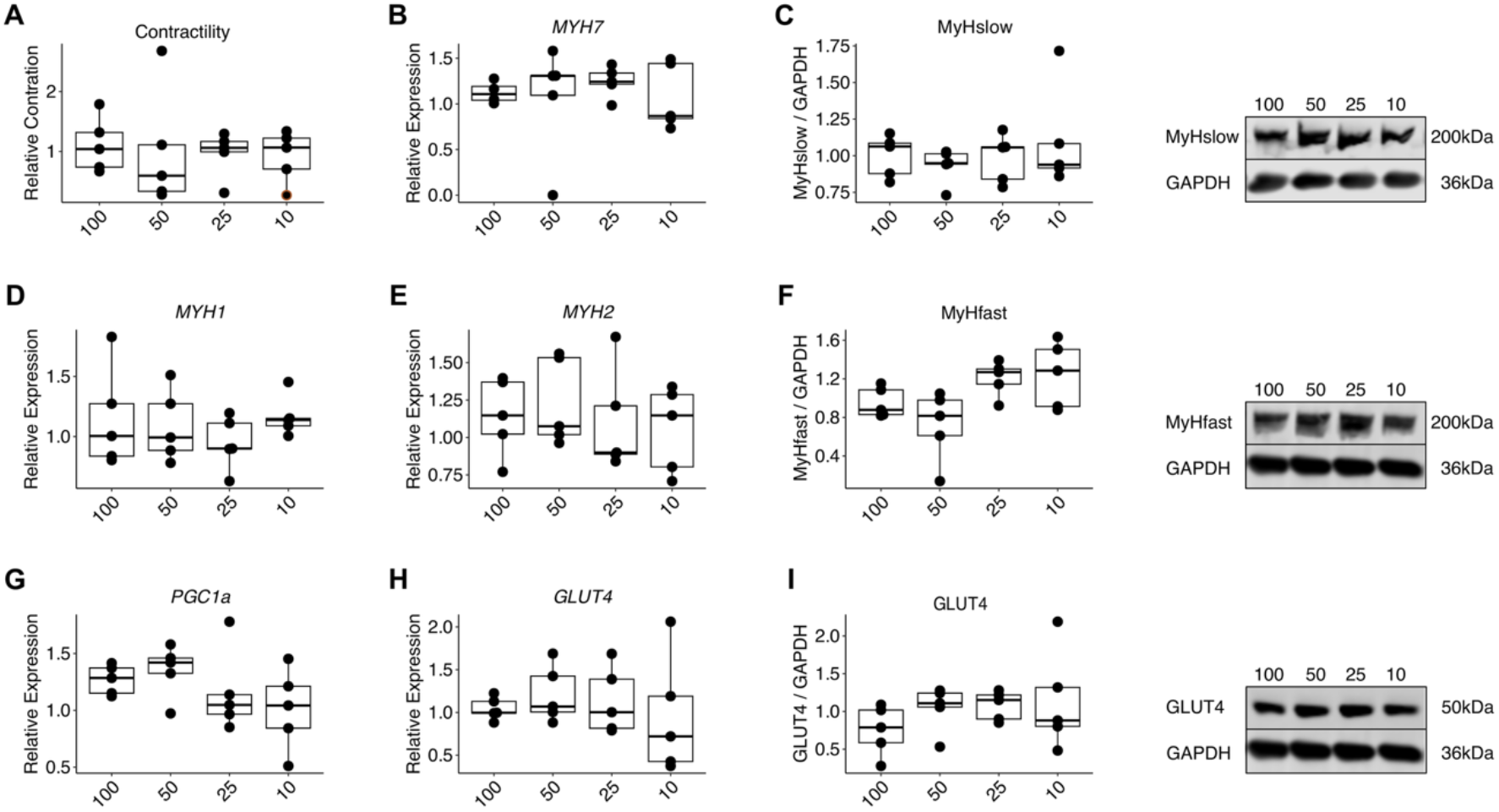
Dose titration of IGF1 during myotube differentiation. Primary human myotubes were differentiated for 8 days in the presence of insulin-like growth factor1 (IGF1) to analyze contraction and differentiation. The IGF1 concentration was kept at 10, 25, 50 or 100ng/ml until harvest on day 8. (A) Contractility represented by contraction distance. RNA expression of differentiation markers *MYH7* (B), *MYH1* (D), *MYH2* (E), *PGC1a* (*PPARGC1A*) (G) and *GLUT4* (H) were measured on day 8 via qPCR. Protein levels of MyHslow (MYH7) (C), MyHfast (MYH1/2) (F) and GLUT4 (I) were measured by Western blot. Representative blots are shown. Statistical significance was determined by one-way ANOVA with Bonferroni correction for multiple comparisons, n=4-5 individual donors, *p<0.05, **p<0.01, ***p<0.001 vs 100.

**Fig. S3.**
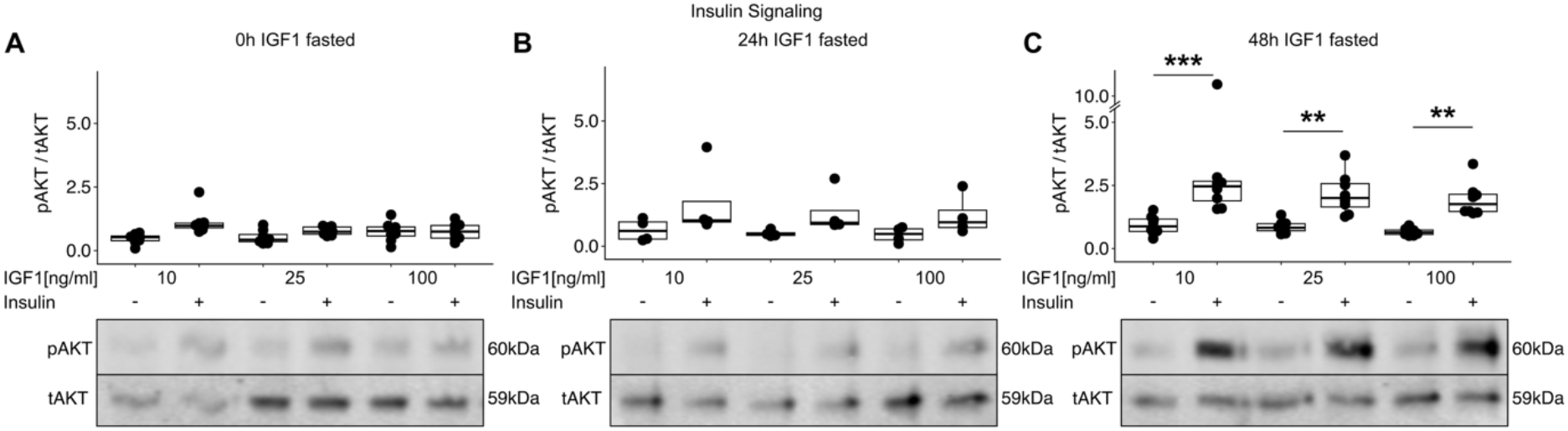
Insulin-dependent AKT phosphorylation. Primary human myotubes were differentiated for 8 days in the presence of 10, 25 or 100ng/ml insulin-like growth factor1 (IGF1) and IGF1 was removed for 0h (A), 24h (B) or 48h (C) before harvest on day 8 with or without prior 10min of 10nM Insulin stimulation. Ser-473 phosphorylation of AKT, and total AKT were measured by Western blot. Representative blots are shown. Statistical significance was determined by one-way ANOVA with Fisheŕs LSD or Welch post hoc test depending on normal distribution, n=4-8 individual donors, **p<0.01, ***p<0.001.

**Fig. S4.**
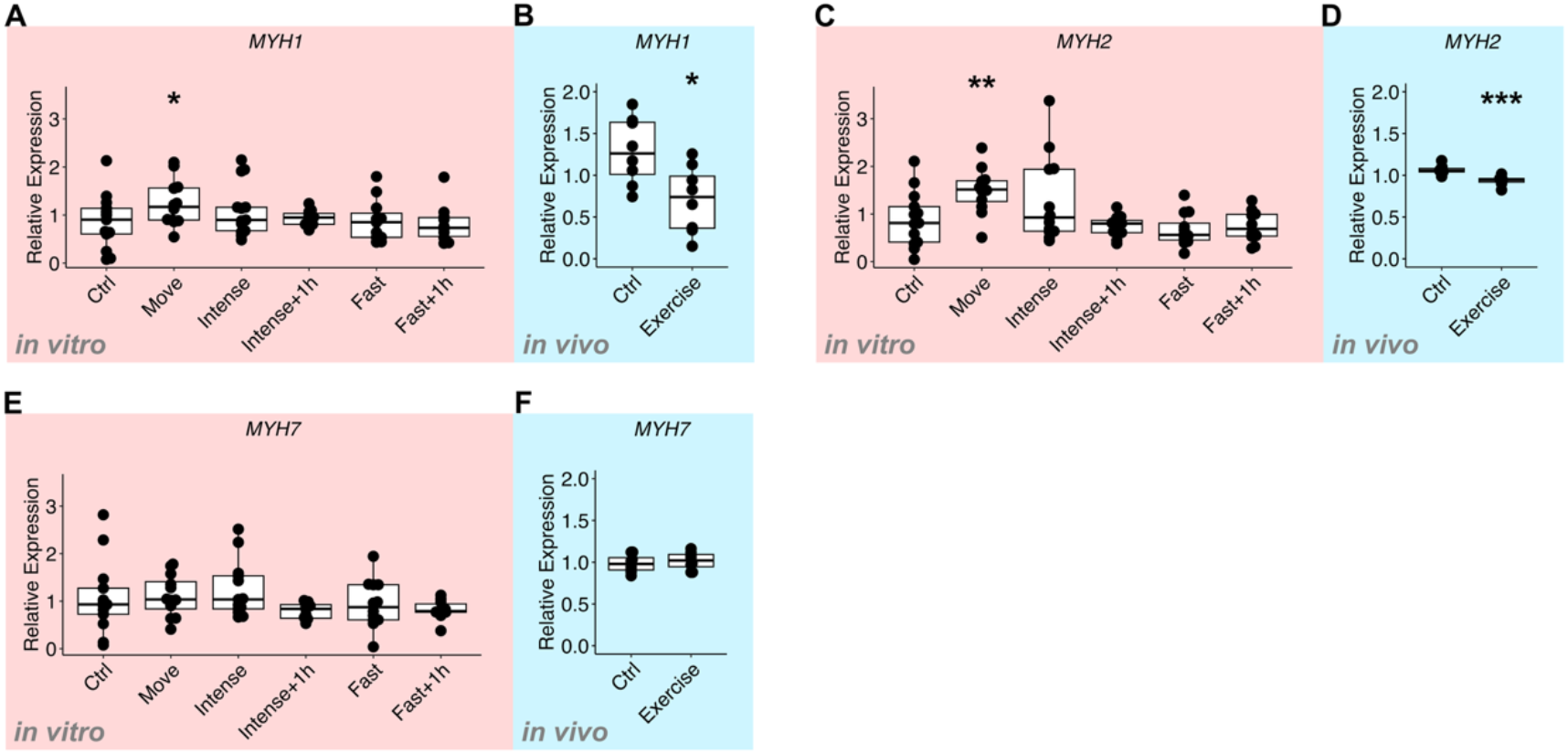
Differentiation marker after in vitro exercise. SMOs were generated from primary human myoblasts and differentiated for 7 days before analysis. RNA expression of *MYH1* (A, B), *MYH2* (C, D) and *MYH7* (E, F) were measured in SMOs directly or 1h (+1h) after in vitro exercise at indicated protocols compared to unexercised controls (Ctrl) (A, C, E) and in skeletal muscle biopsies of the same human donors SMOs were generated from before (Ctrl) and 1h after exercise (Exercise) (B, D, F). Statistical significance was determined by one-way ANOVA with Fisheŕs LSD or Welch post hoc test depending on normal distribution and Bonferroni correction for multiple comparisons when appropriate, n=8-12 individual donors, *p<0.05, **p<0.01, ***p<0.001.

**Fig. S5.**
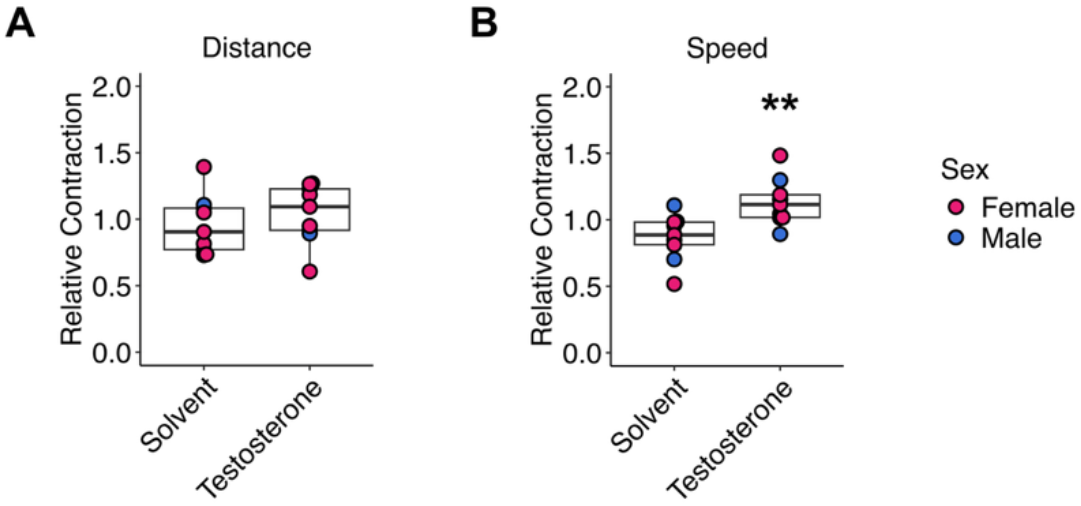
Testosterone modulates human skeletal muscle contractility. SMOs were generated from primary human myoblasts and differentiated for 7 days before analysis Contraction (A) distance and (B) speed of SMOs stimulated with the “Move” protocol were measured after treating SMOs during 7 days of differentiation with or without 100nM testosterone. Donor sex is indicated, pink dots, female, blue dots, male. Statistical significance was determined by one-way ANOVA with Fisheŕs LSD or Welch post hoc test depending on normal distribution, n=10 individual donors, *p<0.05, **p<0.01, ***p<0.001.

